# Comparative Analysis of Transposable Elements in *Hermetia illucens*

**DOI:** 10.64898/2026.07.10.737754

**Authors:** Hector Rosche-Flores, Sean Fischer, Christine J Picard

## Abstract

**Background:** The black soldier fly (*Hermetia illucens*) is an emerging model for bioconversion and industrial rearing. Its genome is highly repetitive, yet the contribution of transposable elements (TEs) to population divergence and demographic processes. The sampled populations represent a gradient of demographic histories, including wild and near-wild North American populations, and domesticated European strains with shared industrial origins. Difference in TE composition may influence genome structure, regulatory variation, and evolutionary responses to captive environments.

**Results:** A comparative analysis of the repetitive landscape was done for four *H. illucens* genomes, one of which is a wild-caught specimen. Total repeat content was high across all assemblies (67.6% to 70.8%) and dominated by LINE elements. Class-level TE diversity was nearly identical among genomes, but multiple DNA transposon families showed distinct lineage-specific differences. Large families including *Maverick* and *Academ* were generally depleted relative to the wild sample. Divergence profiles revealed patterns consistent with recent turnover in several families. Family level turnover, rather than class level change, accounted for the most difference among the genomes. TE-associated structural variants (TESVs) were also not uniformly distributed. Most chromosomes showed mid-chromosome enrichment, and a pronounced TESV peak on chromosome 5 overlapped a histone rich region containing many unclassified repeats. Use of a repeat library derived from multiple genomes increased the number of detected TESVs and improved classification within complex regions, demonstrating that multi-genome libraries enhance annotation accuracy compared to single reference-based models.

**Conclusions:** Multiple DNA transposon families show evidence of recent or lineage-specific amplification in *H. illucens*, suggesting that TE amplification contributes to genome variation during demography-associated TE turnover. The multi-genome-based library improved TE detection and classification, providing a proof of concept that even a small lineage-inclusive repeat library enhances annotation accuracy and capture TE diversity missed by single-reference approaches. Together, these findings demonstrate that TE family turnover plays a significant role in shaping genome architecture and adaptation in this species.

## Background

Transposable elements (TEs) are well known to shape genomes in both structure and evolution [1, 2], influencing gene regulation, chromosomal architecture, and structural variation [3]. Their dynamics are shaped by intrinsic properties such as replication mechanisms as well as extrinsic factors such as population size, selection, and demographic histories [4]. While some organisms, like the model *Drosophila melanogaster*, have simple TE landscapes, many non-model organisms differ significantly and remain challenged with annotation biases and limited lineage-specific resources, which may obscure true TE diversity [5]. Systems that combine high repeat content with well-defined demographic contexts offer a powerful framework to disentangle these effects. The black soldier fly (*Hermetia illucens*, Diptera: Stratiomyidae) represents such a system, with large, TE-rich genomes and populations spanning wild and recently bottlenecked populations, as well as long domesticated lineages [6].

*Hermetia illucens* (black soldier fly, or BSF) is a saprophagous fly whose larvae develop on decomposing organic matter such as plant waste and manure [7]. BSF is regarded as endemic to the Americas, with the earliest confirmed records of BSF outside the Americas date to the early twentieth century, including reports from South Africa in 1914, Malta in 1926, and Hawaii in 1930, with subsequent detections across the Pacific, Africa, Europe, and Asia appearing from the 1930s to 1950s onward [6], however, genetic data suggest global expansion well before anthropogenic forces [8]. BSF larvae exhibit rapid growth and a tolerance of a broad range of organic substrates. Because of these traits, BSF has become a central species in industrial- and garden-scale rearing systems, where it is deployed to valorize food waste and agro-industrial byproducts for use in animal feed [9, 10]. Furthermore, BSF has been explored for potential use in biodiesel production, cosmeceuticals, and antibiotic alternatives for livestock [11–14]. As BSF entered industrial rearing systems, commercially maintained lineages began experiencing population dynamics that differ markedly from those of wild populations, including likely bottlenecks, altered mating structure, and sustained artificial selection. The recent global expansion potentially provides a useful contrast between historical population structure [8] and contemporary, human-mediated demographic spread.

Despite this relatively recent documented history, BSF exhibits a surprisingly deep and complex evolutionary background. Surveys consistently report high global genetic diversity, with deeply divergent mitochondrial and nuclear lineages found across wild and human-associated populations [8, 15–19]. Whole-genome and mitochondrial analyses further reveal major lineages whose estimated divergence times extend into the late Miocene and Pliocene, indicating that present-day populations reflect a far older and more structured demographic history [8]. Together, these findings highlight *H. illucens* as a species with substantial standing genetic variation and genetic patterns that vary strongly across geographic regions, features that shape how modern populations respond to both natural and managed environments.

The process of domestication and its sustained artificial selection compress genetic variation into narrow, stable genomic backgrounds – useful in mass production scenarios. However, this homogenization is known to constrain adaptive potential, alter expression dynamics and facilitate the accumulation of deleterious variants which is already being observed in some long running commercial BSF operations [20–24]. Beyond the effects on coding sequences, such demographic histories are hypothesized to influence the abundance, activity, and regulation of transposable elements (TEs). TEs are ubiquitous components to eukaryotic genomes and can follow diverse evolutionary trajectories depending on host or demographic contexts. TEs may proliferate rapidly, become epigenetically silenced, or be co-opted into regulatory or coding functions [25]. Hosts in turn deploy multiple defense mechanisms, including small RNA-mediated silencing, DNA methylation, and heterochromatin formation, to suppress deleterious TE activity, and mismatches between TE load and host control systems can result in severe fitness consequences [1, 26, 27].

The total repeat content found in Insect species varies dramatically, reflecting lineage-specific histories of transposable element expansion and loss. A comparative analysis of 73 arthropod genomes found extensive variation of TE content, recent activity, and superfamily diversity across and within insect orders as well as large fractions of unclassified repeats [28]. Across Diptera, TE content spans from <5% to >60% of the genome structure, with much variation among orders, and even within genera [29]. While *Drosophila melanogaster* was instrumental in discovering much of what we know about TEs [30], its genome is compact compared to *H. illucens* [31] and is atypical of many non-model dipterans [32]. One study reported genome sizes between 174 and 253 Mbp and that these differences, as well as those for other Drosophila species, were correlated with TE content [33]. By contrast, many other insect lineages exhibit much higher proportions of repetitive DNA, with *Hermetia illucens* assemblies exceeding 65% [5].

The expansion of TEs in arthropods appears to be episodic bursts of activity within specific TE families, followed by periods of relative inactivity [28, 34]. Comparable burst-like dynamics of TE proliferation have been reported across a wide range of taxa, including mammals, bats, fish, and several plant lineages [35]. However, comparisons of repeat content across taxa are complicated by annotation bias. Most repeat annotation pipelines rely on reference databases such as RepBase and Dfam, both heavily curated from Drosophila species. As a result, the proportion of unclassified repeats increases with genetic distance from Drosophila, rising from roughly 13% in Drosophila species to over 40% on average in other insect orders and exceeding 70-85% in early diverging and undersampled groups such as Thysanoptera (thrips) and Ephemeroptera (mayflies) [5]. Because TE families undergo rapid lineage-specific turnover, reference libraries derived from distant taxa have limited utility, as demonstrated by large-scale drosophilid surveys [36].

Repetitive DNA is also frequently underrepresented in the genome assemblies themselves, as collapsed or filtered repeat rich regions can obscure true TE abundance and diversity [37]. *De novo* discovery in non-model insects is further complicated by assembly quality: analyses of Stratiomyidae genomes show a slight negative correlation between assembly *BUSCO* completeness and proportion of repeat content identified [38]. Together, this highlights the need for a more comprehensive, diverse lineage-specific repeat library to accurately characterize the repeat landscape of *H. illucens*. Beyond improving annotation accuracy, uncovering lineage-specific and previously unclassified repeats in *H. illucens* provides an opportunity to examine how repetitive DNA contributes to genome evolution under domestication. Differences in TE abundance or recent activity may reveal the genomic consequences of population bottlenecks, altered selection regimes, or natural adaptations to captive environments.

The populations examined here represent a gradient of demographic histories, including a wild North American population, a near-wild population subject to recent bottlenecks, and two long domesticated strains with shared industrial origins. Although most of them ultimately trace back to North America, they differ substantially in their recent demographic trajectories, providing a framework to study the impact of these conditions on TE landscapes in *Hermetia illucens*. The aim of this work is to characterize and compare the transposable element landscape of wild and domesticated black soldier fly lines, assessing differences in total TE content, class diversity, and signatures of recent activity (e.g. sequence divergence-based proxies) to identify patterns that may reflect demographic history or adaptation to captive environments.

## Results

### Genome assemblies

We generated two *de novo Hermetia illucens* genome assemblies and analyzed two additional assemblies from external sources (Table 1). A genome assembly from a wild specimen collected in Indianapolis, Indiana, USA (IN) was produced from PacBio HiFi data totaling 16.49 Gb (≈16× coverage). The IN assembly had a total length of 1.02 Gb across 1,443 contigs with a contig N50 of 1.71 Mb and Diptera *BUSCO* completeness of 90.0%. A second *de novo* assembly (CA) was generated from a specimen collected in California from a population generated from wild samples and maintained in captivity for approximately five generations prior to sequencing. The CA assembly had a total length of 1.20 Gb across 18,393 contigs with a contig N50 of 0.40 Mb and Diptera *BUSCO* completeness of 94%. Two additional *H. illucens* assemblies were obtained from external sources. The UK (reference) assembly downloaded from NCBI had a total length of 1.01 Gb and consisted of 21 contigs including mitochondrial and unplaced scaffolds; seven contigs corresponded to near chromosome-scale sequences, and the assembly had a contig N50 of 180.46 Mb with Diptera *BUSCO* completeness of 95.6%. The IT assembly provided by the authors had a total length of 0.89 Gb across 169 contigs with a contig N50 of 162.19 Mb and Diptera *BUSCO* completeness of 84.7%. While these assemblies differ substantially in contiguity and sequencing technology, downstream analyses were designed to focus on relative TE composition and divergence patterns that are robust to assembly fragmentation (see Methods).

**Table 1.**
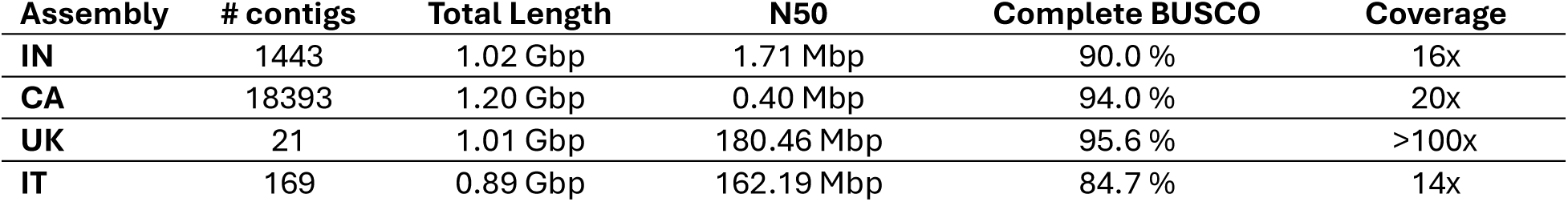
Summary statistics for genome assemblies.

### TE class composition across genomes

Total repeat content was high across all four genomes, ranging from 67.6% to 70.8% (Figure 1). These values are consistent with previous estimates [5], and the slightly higher repeat fractions observed here may reflect improved detection using a custom repeat library derived from all four genomes. Across all assemblies, LINE elements dominated, accounting for ∼40-50% of the genome, while DNA transposons and LTR elements contributed smaller and broadly comparable fractions (Figure 1). The IN genome contained the lowest total repeat proportion, but the relative abundance of the major classes was otherwise consistent among assemblies.

**Figure 1.**
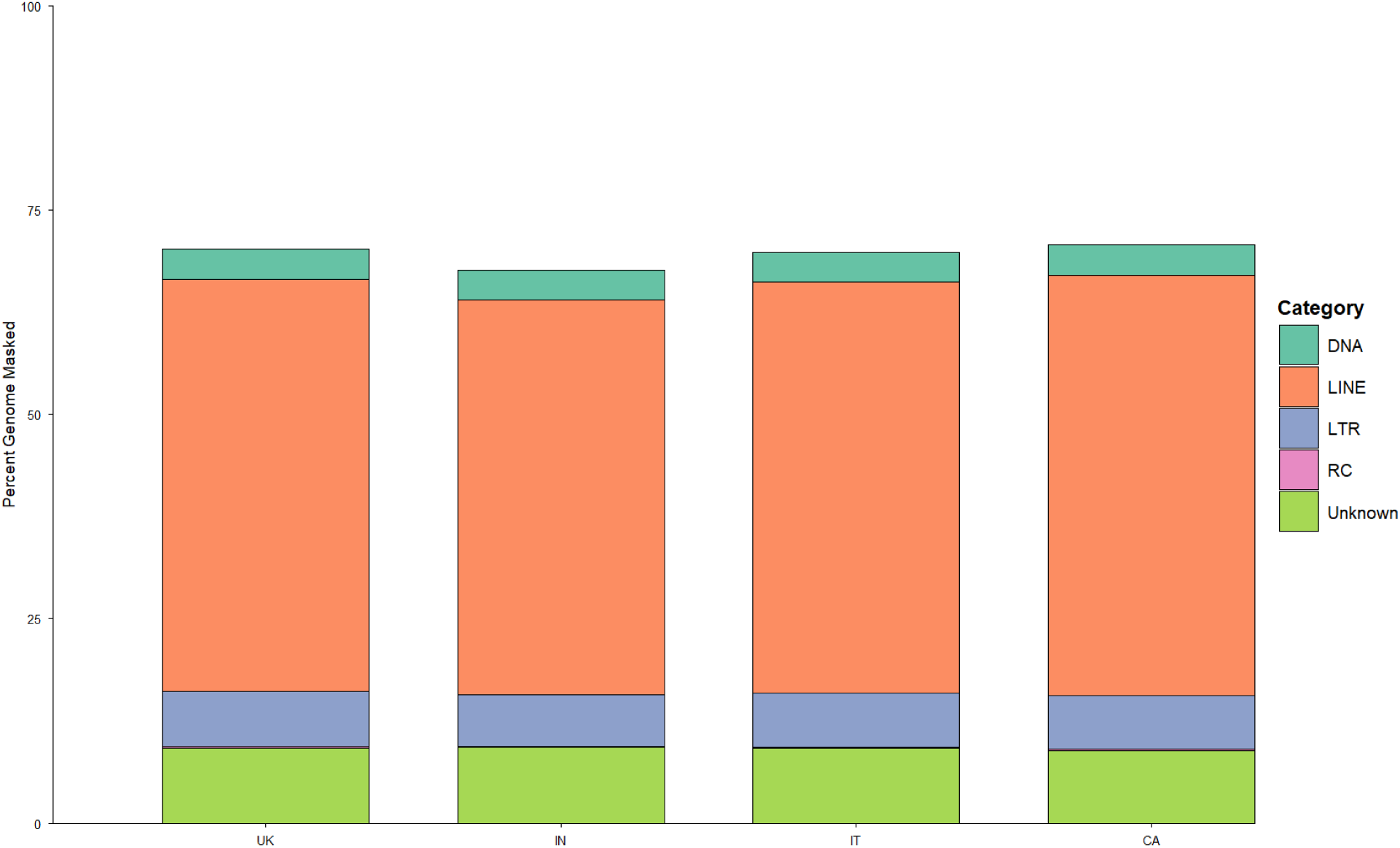
Genome-wide repeat-masked content for each assembly, showing the proportion of transposable element (TE) orders: DNA transposons (DNA), long interspersed nuclear elements (LINE), long terminal repeat elements (LTR), rolling-circle elements (RC), and unclassified/unknown repeats.

### Superfamily-level shifts in domesticated genomes

To quantify superfamily-level changes associated with domesticated lineages, we compared relative superfamily abundance in CA, IT, and UK to the wild IN genome (Figure 2). The non-LTR superfamily R2 showed the largest divergence among superfamilies, with a dramatic increase in IT (+ 214.1 %), a modest enrichment in CA (+ 21.6 %), and a slight reduction in UK (- 14.0 %). Among DNA transposons, *Academ* showed the largest and most variable reductions (from -16.2 % to -63.0 %). The *Maverick* superfamily also declined across all domesticated genomes (-14.5 % to -42.2 %). In contrast, *Sola* increased in abundance in every domesticated genome with modest enrichment in CA (6.9 %)(+6.9%), and substantially larger increases in longer-domesticated IT and UK genomes (60.2 % and 51.3 %, respectively).

**Figure 2:**
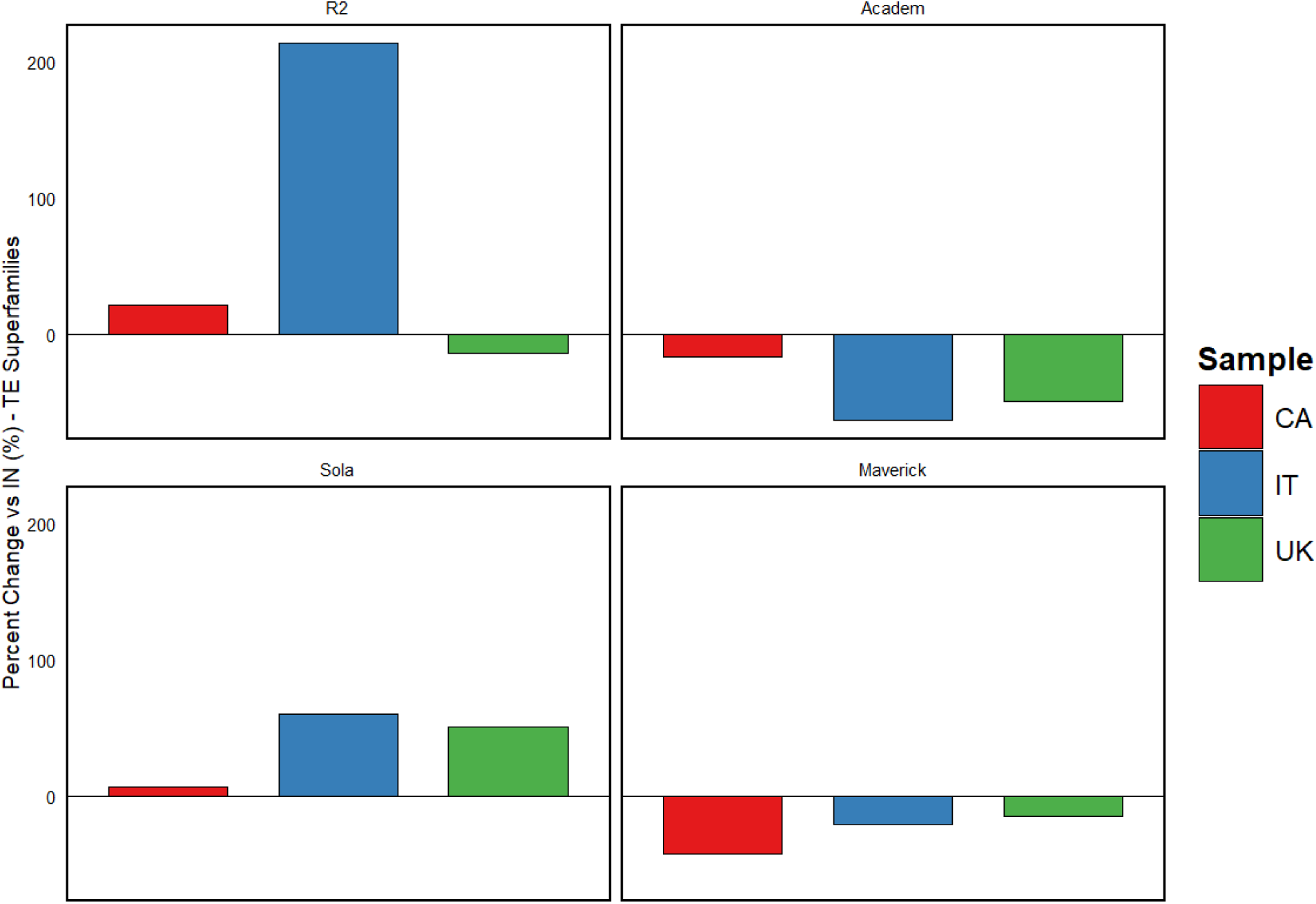
Differences in major TE superfamily abundance relative to the wild IN genome for domesticated genomes CA, IT, and UK.

### TE divergence profiles indicate recent activity

Using sequence divergence from the consensus as a proxy for relative age estimates to compare the distribution of more recent versus ancient TE copies across order (Figure 3). Unsurprisingly, across genomes, 16-44 % of the total TE content had <5 % sequence divergence from the consensus, consistent with a fraction of the TE elements being relatively young, however the proportion of low divergent copies varied by TE order. Rolling circle (RC) elements occupied the largest fraction of recent copies (38% - 44 %), whereas DNA transposons and LTR elements were intermediate (DNA: 24 – 27 %; LTR: 20 – 23 %). In contrast, LINE elements contain 16 -17 % low divergence copies and were dominated by older insertions (67-68 %), while unclassified repeats were almost all ancient (∼82-84 %).

**Figure 3.**
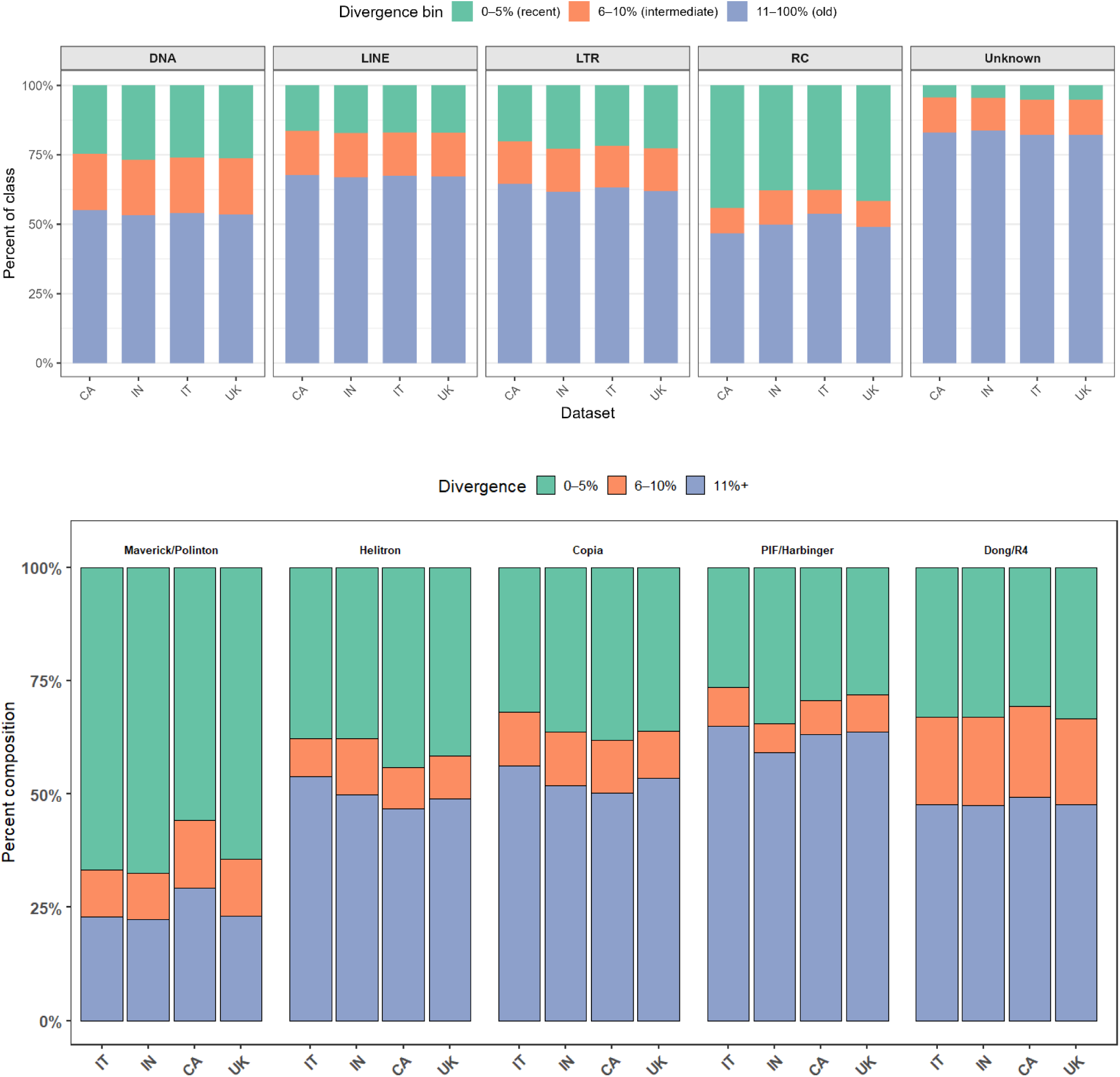
Percentage of each genome with sequence divergences across TE orders (a) and superfamilies (b).

At the superfamily level, several lineages showed substantial fractions of low divergence copies (Figure 3). *Copia* elements showed 28-32 % of copies in the 0-5% divergence class across genomes. The LINE superfamily *Dong/R4* exhibited similar levels (31-33%), with little variation among wild and domesticated lineages. *Helitron* (RC) elements exhibited the highest proportion of young insertions, ranging from 38% in the wild genome to 44% in CA. DNA transposon superfamilies showed more variable profiles: *Maverick* elements contained the largest fraction of low-divergence copies (56-67 %) while *PIF/Harbinger* elements showed more modest levels (26-35 %). Collectively, these results indicate that recent TE activity in *H. illucens* is contributed by multiple superfamilies, with *Helitron*, *Copia*, *Dong/R4*, and especially *Maverick* elements representing prominent sources of ongoing turnover.

### Genomic distribution of focal TE superfamilies relative to genes (UK reference)

To examine the genomic context of focal TE superfamilies, we annotated TE positions in the UK reference genome (Table 2). TE insertions were classified as exonic, intragenic, proximal (within 2kbp of a gene), or intergenic. Total copy numbers varied widely across superfamilies, ranging from 143 (*Academ*) to 58,383 (*Dong/R4*), with intermediate counts for *Copia* (1,416) *PIF/Harbinger* (1442), *Maverick* (1,672), *Helitron* (3,971), and *Sola* (383).

**Table 2.** Distribution of TE superfamilies in the UK reference genome by genomic feature.

| **Superfamily** | **Total** | **Exonic** | **Intragenic** | **Proximal** | **Intergenic** |
| --- | --- | --- | --- | --- | --- |
| *Academ* | 143 | 1 | 54 | 9 | 79 |
| *Copia* | 1416 | 7 | 816 | 56 | 537 |
| *Dong/R4* | 58383 | 161 | 28527 | 3151 | 26544 |
| *Helitron* | 3971 | 68 | 2094 | 238 | 1571 |
| *Maverick/Polinton* | 1672 | 96 | 403 | 91 | 1082 |
| *PIF/Harbinger* | 1442 | 71 | 759 | 112 | 500 |
| *Sola* | 383 | 1 | 190 | 28 | 164 |

Across superfamilies, insertions were nearly evenly divided between gene-associated regions (exonic, intragenic, and proximal) and intergenic regions. Superfamilies varied in bias: *Copia*, *Helitron*, *PIF/Harbinger*, and *Sola* showed enrichment in gene-associated regions (51-65%), whereas *Maverick* and *Academ* were biased towards intergenic (55-65%). *Dong/R4* was evenly distributed across gene-associated and intergenic contexts.

### Functional enrichment of *Sola*-associated genes

Among the superfamilies examined, *Sola* was consistently enriched in all three domesticated genomes relative to the wild IN assembly and showed a modest bias towards insertion within or near genes. To test whether *Sola* insertions were associated with genes involved in particular biological processes, we compared the gene ontology (GO) annotation of *Sola*-associated genes with those genes intersecting any other *RepeatMasker*-classified TE superfamily. Collapsing fine-scale GO terms to parent categories reduced the effective number of statistical tests, yielding in a robust set of enriched biological processes after Benjamini–Hochberg correction (Figure 4).

**Figure 4.**
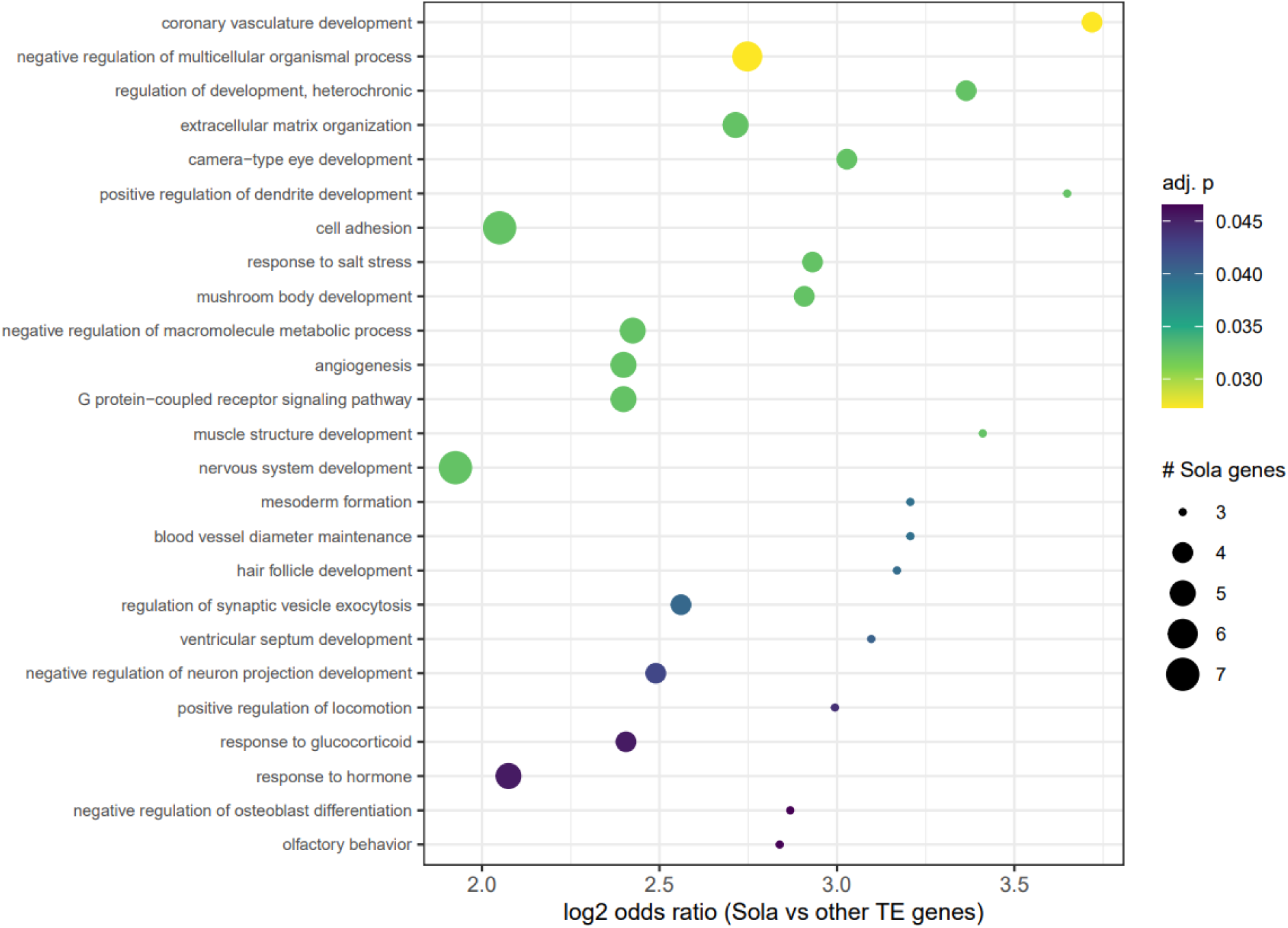
Functional enrichment of genes associated with Sola transposable element insertions.

*Sola* insertions were significantly overrepresented in genes associated with developmental regulation, including regulation of development, heterochronic (OR = 10.3; p_adj_ = 0.0325) and negative regulation of multicellular organismal process (OR = 6.7; p_adj_ = 0.0273). Enriched categories also included several tissue and organ morphogenesis terms, such as coronary vasculature development (OR = 13.2; p_adj_ = 0.0273), mesoderm formation (OR = 9.23; p_adj_ = 0.0395), camera-type eye development (OR = 8.16; p_adj_ = 0.0325), and muscle structure development (OR = 10.6; p_adj_ = 0.0325). GO enrichment also identified significant overrepresentation of categories related to neural and behavioral processes including positive regulation of dendrite development (OR = 12.5; p_adj_ = 0.0325), nervous system development (OR = 3.8; p_adj_ = 0.0325), olfactory behavior (OR = 7.15; p_adj_ = 0.0465), and regulation of synaptic vesicle exocytosis (OR = 5.9; p_adj_ = 0.0400). Additional significant terms included responses to hormone (OR = 4.21; p_adj_ = 0.0454), negative regulation of macromolecule metabolic process (OR = 5.37; p_adj_ = 0.0325), and G protein coupled receptor signaling (OR = 5.27; p_adj_ = 0.0325).

### Diversity of TEs by Order and Superfamily

Shannon diversity and Pielou’s evenness were broadly consistent across assemblies at both Order and Superfamily levels, indicating no substantial differences in overall TE diversity, although the UK sample showed slightly lower Order level diversity and higher family level evenness (Supplementary X).

### Chromosomal distribution of TE-associated structural variants (TESVs)

To explore how recent TE activity relates to structural variation, we examined the chromosomal distribution of TE-associated structural variants (TESVs) (Figure 5). Most chromosomes showed broadly similar TESV profiles, with variant density highest near the putative centromeres of chromosomes 1-4, whereas chromosomes 5 and 6 show more evenly and reduced density across its chromosomes.

**Figure 5:**
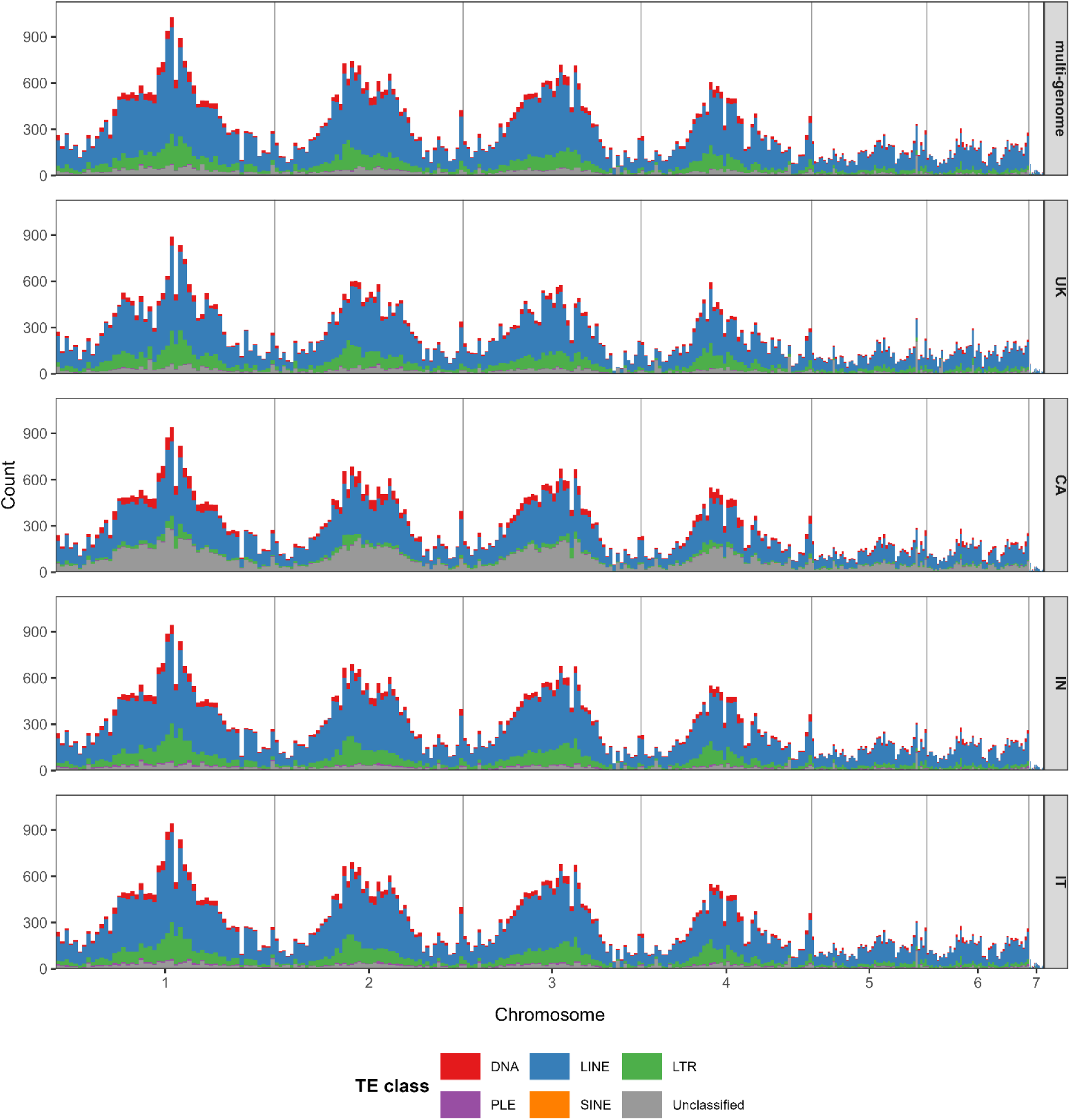
Distribution of transposable element structural variants (TESVs) across chromosomes relative to the reference UK genome.

## Discussion

### Repeat landscape stability and class-level conservation

Total repeat content across *Hermetia illucens* genomes ranged from 67.6 % to 70.8 %, with the reference assembly showing 70.2 % masked bases. These values are relatively high for Diptera but broadly consistent with previous estimates and with emerging patterns from repeat-rich non-model insects[5]. The slightly elevated repeat fractions reported here may reflect improved detection enabled by the multi-genome-derived repeat library, which is expected to capture lineage-specific diversity that may be underrepresented in single-reference approaches. Despite modest variation in total repeat abundance, the proportional composition of the major TE classes was conserved across genomes including a ∼3% lower repeat fraction in the wild IN genome. LINE elements consistently accounted for approximately 40-50 % of total repeats, while DNA transposons and LTR retrotransposons contributed smaller and broadly comparable fractions. Together, these results indicate that repeat-rich genome architecture is broadly conserved at the class level across populations with differing demographic histories, suggesting that large-scale TE class composition is relatively insensitive to recent population processes.

### Lineage-specific superfamily responses

In contrast to class-level conservation, we observe lineage-specific shifts at the superfamily level. The non-LTR retrotransposon superfamily R2 exhibited the greatest difference among all superfamilies examined, with variable abundance across genomes, including enrichment in CA and IT and a reduction in the UK genome. Given that R2 elements are typically associated with rDNA loci, this pattern may reflect lineage-specific differences in rDNA copy number or turnover, although this cannot be directly resolved with the present data [39].

Among DNA transposons, *Academ* and *Maverick* superfamilies both declined in domesticated genomes but followed distinct trajectories. This pattern is consistent with expectations under repeated bottlenecks and altered selective regimes [40], although alternative explanations (e.g. lineage-specific history or assembly effects) cannot be excluded. In contrast, *Maverick* elements showed their strongest depletion in CA, with more moderate reductions in IT and UK suggesting that lineage-specific demographic history, rather than cumulative domestication time alone, may contribute to the observed differences.

*Sola* elements showed the opposite pattern, with consistent enrichment across all domesticated genomes. However, the magnitude of enrichment differed strongly among lineages, with only a modest increase in CA but much larger increases in IT and UK. Although *Sola* enrichment is consistent across the domesticated genomes, the underlying mechanisms (e.g. transposition, excision or differential retention) cannot be distinguished here. Together, these results indicate that demographic histories under captive expansions reshape TE composition in *H. illucens* through heterogeneous, superfamily-specific responses rather than uniform expansion or loss.

### Divergence profiles reveal heterogeneous TE turnover dynamics

Divergence-based analyses provide insight into the temporal structure of the *H. illucens* repeat landscape. At the order level, a substantial fraction of total TE content consisted of relatively low-divergence copies, indicating the presence of comparatively younger insertions across multiple TE orders. However, the distribution of low divergence copies differed markedly among orders, suggesting that replication strategy and removal dynamics strongly shape the age structure of the repeat landscape.

Rolling-circle elements exhibited the highest proportion of low-divergence copies, suggesting their contribute substantially to ongoing TE turnover within this comparative framework [41]. This pattern is consistent with their strand-displacement replication mechanism, which can generate multiple copies from a single event [42]. The functional consequences of this pattern remain unresolved, and these data do not directly establish links to phenotypic variation or selection, but they may provide a reservoir of structural variation available for selection in captive environments.

DNA transposons and LTR retrotransposons exhibited intermediate divergence profiles, consistent with a balance between insertion and removal processes [2, 43]. By contrast, LINE elements showed the lowest proportion of low-divergence copies and were dominated by older insertions, consistent with long-term accumulation of fragmented remnants and limited recent mobilization. At the superfamily level, recent TE turnover was distributed across multiple lineages rather than dominated by a single superfamily. Several superfamilies consistently maintained substantial fractions of the low-divergence copies across both wild and domesticated genomes. However, the magnitude and consistency of these fractions varied, indicating heterogeneous turnover dynamics. Together, these patterns indicate that TE turnover in *H. illucens* is distributed across multiple lineages, with no single TE order dominating low divergence insertions across all genomes.

### Reconciling *Maverick* abundance and divergence

*Maverick* elements (also known as Polintons) illustrate the importance of interpreting abundance and divergence metrics on different timescales. Although *Maverick* elements exhibited the highest proportion of low-divergence copies across all genomes, they were nonetheless depleted in domesticated genomes. This apparent contradiction highlights that divergence-based metrics do not directly correspond to recent demographic timescales, and low divergence classes likely predate recent domestications [18].

*Mavericks* are large virus-like DNA transposons whose abundance often fluctuates through rare but pronounced bursts of amplification followed by long periods of quiescence and erosion [44–46]. Under this framework, the observed depletion of *Maverick* elements in captive genomes is consistent with demographic processes associated with domestication, such as bottlenecks, founder effects, and rapid expansions that disproportionately reduce large, inactive elements. In the absence of these recent demographic disruptions, *Maverick* copy number might be expected to remain stable or increase over longer evolutionary timescale, but the short and intense history of anthropogenic forces acting on the *H. illucens* likely truncated this trajectory.

### Genomic context of recent TE insertions and functional associations

Across all superfamilies, TE insertions were nearly evenly divided between gene-associated and intergenic regions, indicating that TE turnover in *H. illucens* is not confined to gene-poor genomic compartments. This balance is consistent with expectations for genomes harboring multiple TE lineages with distinct insertion preferences and fitness effects. Individual superfamilies exhibited biases aligned with their structural properties. *Maverick* and *Academ* elements were enriched in intergenic regions, potentially reflecting disruptive potential of large, structurally complex insertions within genes. Both superfamilies were also depleted in captive genomes, raising the possibility that domestication-associated demographic processes have further reduced their persistence in gene-associated regions. In contrast, *Copia*, *Helitron*, *Dong/R4*, *PIF/Harbinger*, and *Sola* elements showed modest enrichment in gene-associated regions. Among these, *Sola* elements were notable in that gene-associated enrichment coincided with increased abundance in captive genomes.

Functional enrichment analyses revealed that *Sola* insertions were disproportionately associated with genes involved in developmental regulation, tissue and organ morphogenesis, neural development, and hormone-mediated signaling. Although enriched GO terms spanned diverse processes, they converged on two biological themes: the regulation of growth and structural development, and the coordination of neural and behavioral responses. These pathways are central to traits commonly targeted, whether directly or indirectly, by artificial selection in commercial rearing environments, including growth rate, body size, stress tolerance and behavior.

While the present analysis cannot establish a causal role for *Sola* insertions for any specific trait, the enrichment of *Sola* associated genes within these pathways raises the possibility shifts in selection pressures under captive propagation may have disproportionately retained *Sola* insertions near genes central to development and environmental responsiveness.

### Chromosomal distribution of TE-associated structural variation

TE-associated structural variants (TESVs) were broadly enriched toward chromosome centers, consistent with the documented reciprocal relationship between recombination rate and TE accumulation. Insertions tend to accumulate in low-recombination regions, and high TE density can further suppress recombination, promoting local structural divergence [47, 48].

Chromosomes 5 and 6, which are substantially smaller than chromosomes 1-4, exhibited distinct TESV profiles. Comparative analyses across insects and plants indicate that smaller chromosomes often experience higher recombination rates per megabase, enabling more efficient purifying selection and limited TE accumulation outside pericentromeric regions [49]. This pattern may explain the more uniform TESV distributions observed for these chromosomes.

A pronounced TESV peak observed on chromosome 5 overlapped a previously identified putative domestication locus characterized by long runs of homozygosity [6]. This region also contained a high fraction of unclassified repeats, even when using a multi-genome derived repeat library. The overlap with tandemly repeated histone genes suggests locally complex genomic architecture, rather than missing library representation, contributes to both elevated TESV density and classification difficulty in this region. The region overlaps a dense array of tandemly repeated histone genes in the reference genome, indicating that part of the unclassified content may correspond to repetitive genic DNA or interspersed satellite-like sequences that mask transposable element identity. These complex regions are often difficult to assemble or classify and may contribute to the elevated TESV density observed near the previously reported putative domestication locus [50, 51].

### Limitations and future directions

Several factors complicate the inference of phenotype-TE relationships. Lineage⍰specific TE activation, horizontal transfer, and demographic history can confound ancestry⍰based interpretations. Moreover, many TE⍰mediated effects act indirectly by altering gene regulation rather than coding sequence, making them difficult to detect using standard homology⍰based annotation frameworks. The black soldier fly exhibits exceptional global genetic diversity, with population level divergence approaching 5% based on microsatellite and whole genome data [18]. Captive breeding programs have also produced measurable genomic divergence in fewer than five years of selective breeding [52].

Given this species’ genetic backdrop, variation in repeat sequence among populations could further contribute to classification uncertainty, as rapidly evolving TEs may diverge beyond recognition by consensus models. Although a high-quality *H. illucens* reference genome is available, current annotations remain largely limited to protein coding features. Improving resolution of TE function will require enriched annotation resources that integrate regulatory features such as promoters, enhancers, and untranslated regions, as well as expression data. Expanding repeat libraries to include additional global populations will further improve classification accuracy and help distinguish genuinely novel repeat lineages from complex or genic repetitive DNA.

## 5. Conclusion

Together, these findings demonstrate that early domestication in *H. illucens* has not fundamentally altered the class-level architecture of its repeat-rich genome, but has instead driven pronounced, lineage-specific turnover among individual transposable element (TE) families. Variation in TE abundance and divergence patterns indicates that recent turnover is concentrated within a subset of DNA transposon superfamilies, most notably *Maverick*/*Polinton* and *Sola*, rather than reflecting uniform change across TE classes.

The contrasting patterns of depletion, persistence, and apparent reactivation observed across TE families are consistent with the effects of repeated bottlenecks, founder events, and altered selective regimes associated with captive propagation. At the chromosomal scale, the non-uniform distribution of TE-associated structural variants further emphasizes the importance of the recombination landscape and local genome architecture in mediating TE accumulation and retention.

Collectively, this work supports a model in which genome evolution under early domestication proceeds through selective filtering and family-level reshaping of standing TE diversity, rather than widespread mobilization or collapse of the repeat landscape. By integrating multiple assemblies and constructing a population-inclusive repeat library, this study highlights both the biological and methodological importance of lineage-aware TE annotation for understanding genome evolution in repeat-rich, non-model species.

## Methods

### Genome sequencing, assembly, and quality assessment

#### Wild Indiana genome (IN: PacBio HiFi)

A *de novo Hermetia illucens* genome was generated from a wild specimen collected in Indianapolis, Indiana, USA using PacBio HiFi sequencing. A PacBio HiFi library with the SMRTbell Prep Kit v3 and PippinHT size selection (insert ∼9.4kb) was sequenced on a PacBio Revio instrument using one SMRT Cell 25M. Reads were assembled with *Hifiasm* using default parameters, dual-scaffold mode, and an estimated haploid genome size of 1.01 Gb [53].

Sequencing produced 1.5 million HiFi reads totaling 16.49 Gb (∼16x coverage). Read quality and length were summarized by the median read quality (Q37) and read length N50 (11.2 Kb). The resulting assembly has a total length of 1.02 Gb across 1,443 contigs with a contig N50 of 1.71Mb Assembly completeness was assessed with *BUSCO* using the Diptera lineage odb10 dataset (complete *BUSCO* = 90 %).

#### Recently captive California genome (CA; Oxford Nanopore)

A second *de novo* assembly was generated from a specimen collected in a California population maintained in captivity for <10 generations prior to sequencing. Nanopore libraries were prepared with the SQK-NBD114-24 kit and sequenced on a PromethION FLO-PRO114M flow cell. Reads were quality-checked using *Nanoplot* v1.42.0 and assembled de novo with *FLYE* v2.9.5 with the parameters –scaffold –nano-raw -g 1.01g [54, 55]. Assembly quality was assessed with *Quast* v5.2.0 and *BUSCO* v5 (Diptera_odb10) [56, 57].Oxford Nanopore sequencing yielded approximately 20X genome coverage, estimated using *Nanoplot* and *FLYE* kmer-based coverage estimators. The resulting assembly had a total length of 1.20 Gb and consisted of 18,393 contigs, with a contig N50 of 0.40 Mb. *BUSCO* completeness using the Diptera odb10 lineage dataset was 94 %.

#### External reference assemblies (UK and IT)

Two additional *H. illucens* genome assemblies were obtained from external sources. The UK assembly was downloaded from NCBI (GCF_905115235.1, termed UK, [6]; total length of 1.01 Gb; 21 contigs including mitochondrial; Diptera *BUSCO* completeness 95.6 %; reported coverage >100×)). The IT assembly was provided directly by the authors (ERS14372541, termed IT [58]; total length 0.89 Gb; 169 contigs; contig N50 162.19 Mb; Diptera *BUSCO* completeness 84.7 %; reported coverage 14×). While these assemblies differ substantially in contiguity and sequencing technology, downstream analyses were designed to focus on relative TE composition and divergence patterns that are robust to assembly fragmentation (see Methods). The UK assembly was used as the reference due to its near chromosome-scale contiguity and high *BUSCO* completeness.

### Transposable element identification and classification

#### Repeat library construction

A custom *Hermetia illucens* transposable element (TE) library was generated using all four genomes through an iterative workflow. *RepeatModeler* v2.0.7 was run sequentially on the assemblies in this order UK → CA → IT → IN to perform de novo identification and preliminary classification of TE consensus sequences [59]. After each iteration, the evolving repeat library was used to mask the next genome using *RepeatMasker* v4.1.9 with the RM_BLAST engine [60]. This approach ensured only previously uncharacterized repeat sequences were incorporated at each step. The resulting four-genome repeat library was used for all genome-wide repeat content estimates, TE diversity calculations, and TE structural variant annotation. In addition, genome-specific *RepeatModeler* libraries were generated for targeted comparisons.

#### TE abundance and divergence analyses

*RepeatMasker* output (.out) files for each genome were parsed using the Perl script *parseRM.pl*. Raw repeat landscapes were subsequently standardized using a custom script sanitize_div_Rfam.py script to normalize output formats and harmonize TE lineage labels according to the unified TE classification system [61, 62]. Genome-wide TE abundance was first summarized at the order level using percent masked bases derived from the sanitized parseRM outputs. Relative abundance differences among genomes were then evaluated at the superfamily level. Given the large number of TE superfamilies identified, downstream analyses focused superfamilies showing the largest deviations in masked content relative to the wild IN genome.

Sequence divergence from consensus was used as a proxy for relative insertion age and recent transpositional activity. Divergence profiles were summarized at both order and superfamily levels by grouping parseRM divergence values into three bins: 0–5% (low divergence), 6–10% (intermediate divergence), and >10% (high divergence). The 0–5% divergence threshold was used to represent relatively recent insertions, consistent with previous comparative genomic studies [64,65].

#### TE Diversity metrics

Diversity statistics were calculated separately for TE orders and superfamilies using the non-redundant masked base counts (nt_masked-minus-double) reported by parseRM.pl. For each genome, these values were treated as abundance estimates. Shannon diversity (H) was calculated as

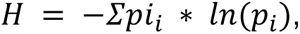

where pi represented the proportional contribution of each TE lineage. Pielou’s evenness (J’) was calculated by normalizing H by the natural logarithm of the number of observed lineages.

#### Genomic context and functional annotation of TE insertions

To assess the genomic distribution of TE insertions, the NCBI-supplied GFF annotation for the UK reference genome was converted to BED format. Non-TE *RepeatMasker* categories (simple repeats, low complexity regions, and structural RNAs) were removed prior to analysis. TE coordinates were intersected with gene models using *BEDTools* [63] and classified as exonic, intronic, proximal to a gene (within 2 Kb of a gene), or intergenic. Gene identifiers associated with overlapping or proximal TE insertions were retrieved from the reference annotation.

Superfamilies showing large abundance differences among genomes were subsequently examined for functional associations. Functional annotations were obtained using *EggNOG-mapper* v2.1.11, in protein mode with default parameters, using the UK proteome as input [64–66]. Gene ontology (GO) enrichment analyses were performed using the clusterprofiler R package, with enrichment statistics normalized by resampling superfamily-associated genes to control differences in gene set size [67].

### TE structural analysis

Transposable element-associated structural variants (TESVs) were identified using the assembly-based variant detection and annotation framework *GraffiTE* [68]. Genome assemblies were aligned to the UK genome and *GraffiTE* generated a multi-sample multi-genome representation from these alignments. Structural insertions and deletions were called using its integrated SV-calling engine (svim-asm) for whole-genome alignments and annotated against the supplied repeat library [69]. Because this analysis focused on TE presence/absence rather than allelic genotyping, only the annotated, multi-genome variant call format (VCF) file produced after *GraffiTE*’s SV calling and TE annotation was used. *GraffiTE* was run separately under five repeat library configurations to evaluate the effect of library composition on TE detection and classification: four runs using single-genome *RepeatModeler*2 libraries and one using the custom multi-genome library. Unified VCF files were imported into R using the vcfr package [70], and TESVs distributions were visualized as stacked density plots in R.

## Availability of supporting data

The UK genome assembly was obtained from NCBI (GCF_905115235.1). The IT genome assembly is available under BioProject PRJEB58627. The two *de novo* assemblies generated in this study are available under BioProject XX.

## Additional files

Custom scripts used in the generation of this publication can be viewed in the github repository found at https://github.com/hroscheflores/myproject

## Competing interests

The authors declare that they have no competing interests.

## Authors’s contributions

H.R.F. and C.J.P. contributed to conceptualization of the study. Data analysis was performed by H.R.F. and S.A.F. The manuscript was written by H.R.F. and C.J.P. All authors reviewed and approved the final version of the manuscript.

## Acknowledgements

The authors would like to acknowledge the support of the Industry Advisory Board of the NSF Center for Insect Biomanufacturing and Innovation (CIBI) for thoughtful conversations and sample acquisitions. This material is based upon work supported by the National Science Foundation under Grant Nos. IIP-2052565, 2052454 and 2057288.

## Notes

https://github.com/hroscheflores/myproject

### Competing Interest Statement

The authors have declared no competing interest.

