## Supplemental Table 1 for "Comparative Analysis of Transposable Elements in *Hermetia illucens*"

**Supplemental Table 1.** Per genome Shannon diversity statistics of transposable elements by TE class and family.

| Sample | Class Shannon | Family Shannon | Class Pielou | Family Pielou |
|---|---|---|---|---|
| IN | 0.898979271 | 2.365591 | 0.558567 | 0.614416 |
| CA | 0.879147279 | 2.360837 | 0.546245 | 0.613181 |
| UK | 0.896587997 | 2.367145 | 0.557081 | 0.614819 |
| IT | 0.891107426 | 2.360948 | 0.553676 | 0.61321 |

At the class level, all non-ref assemblies show nearly identical Shannon diversity (0.995–0.999) and evenness (0.511–0.513), indicating a consistent proportional representation of major TE classes across both wild-caught and captive *H. illucens* populations (Table 1). The limited number of TE classes means that the loss or expansion of any single class would drastically distort diversity values, but this is not apparent among the wild or research colony assemblies.

At the family level, diversity is higher across all assemblies (2.12-2.35), reflecting the larger number of family-level classifications. The reference genome exhibits the highest family level Shannon index (2.347) and evenness (0.645), while wild and recently captive populations show slightly lower but similar values (~2.12; 0.565 respectively). Among the wild populations, Indiana and California samples are indistinguishable in TE diversity metrics despite the California colony having been maintained in captivity for several generations at the time of sequencing. Together, these results indicate that TE diversity and evenness are broadly conserved across the *H. illucens* lineages, with only minor deviation in the long-term laboratory strain. These values are comparable to those observed across large-scale mammalian TE surveys where class level Shannon indices typically range between 0.6 and 1.5, suggesting that the observed reduction in the reference genome falls within falls within the expected natural range [29].
